# Cholesterol promotes both head group visibility and clustering of PI(4,5)P_2_ driving unconventional secretion of Fibroblast Growth Factor 2

**DOI:** 10.1101/2021.04.16.440132

**Authors:** Fabio Lolicato, Roberto Saleppico, Alessandra Griffo, Bianca Pokrandt, Hans-Michael Müller, Helge Ewers, Hendrik Hähl, Jean-Baptiste Fleury, Ralf Seemann, Britta Brügger, Karin Jacobs, Ilpo Vattulainen, Walter Nickel

## Abstract

Fibroblast Growth Factor 2 (FGF2) is a cell survival factor involved in tumor-induced angiogenesis. FGF2 is secreted through an unconventional secretory pathway based upon direct protein translocation across the plasma membrane. Here we demonstrate that both PI(4,5)P_2_-dependent FGF2 recruitment at the inner plasma membrane leaflet and FGF2 membrane translocation into the extracellular space are positively modulated by cholesterol in living cells. We further reveal cholesterol to enhance FGF2 binding to PI(4,5)P_2_-containing lipid bilayers in a fully reconstituted system. Based on extensive atomistic molecular dynamics simulations and membrane tension experiments, we propose cholesterol to modulate FGF2 binding to PI(4,5)P_2_ by (i) increasing head group visibility of PI(4,5)P_2_ on the membrane surface, (ii) increasing avidity by cholesterol-induced clustering of PI(4,5)P_2_ molecules triggering FGF2 oligomerization and (iii) increasing membrane tension facilitating the formation of lipidic membrane pores. Our findings have general implications for phosphoinositide-dependent protein recruitment to membranes and explain the highly selective targeting of FGF2 towards the plasma membrane, the subcellular site of FGF2 membrane translocation during unconventional secretion of FGF2.

## Introduction

Beyond the ER/Golgi dependent secretory pathway through which signal-peptide-containing secretory proteins are transported into the extracellular space (1–4), additional mechanisms of protein secretion evolved in eukaryotic cells. These processes have collectively been termed ‘unconventional protein secretion’ (UPS) (5–7). One of the best characterized UPS cargoes is Fibroblast Growth Factor 2 (FGF2) (6, 8), a potent mitogen involved in fundamental processes of pathophysiological significance such as tumor-induced angiogenesis and the generation of survival signals controlling programmed cell death (9–11). In previous work, FGF2 has been demonstrated to be secreted by a molecular mechanism that is based on direct translocation across the plasma membrane (type I UPS) (5–7, 12–15). Other cargo molecules making use of a type I UPS pathway include HIV-Tat, Tau and, under certain physiological conditions, Interleukin 1β (IL-1β) (5–7, 16, 17), proteins with crucial roles in viral replication, neurodegenerative disorders and inflammatory diseases.

The type I UPS pathway, by which FGF2 is secreted into the extracellular space, is initiated by FGF2 recruitment at the inner plasma membrane leaflet. At this location, FGF2 has been shown to undergo sequential physical interactions with the cytoplasmic domain of the Na,K-ATPase (18, 19), Tec kinase (20–22) and the phosphoinositide PI(4,5)P_2_ (14, 15, 22–24). While the specific functions of the Na,K-ATPase and Tec kinase in this process are only beginning to emerge (5, 6, 19–22), the role of PI(4,5)P_2_ is understood in great detail. In the initial step, binding to PI(4,5)P_2_ triggers oligomerization of FGF2 (22, 25, 26). This process leads to the formation of a lipidic membrane pore with a toroidal architecture accommodating membrane-spanning FGF2 oligomers (5, 14, 22). Once FGF2 oligomers become accessible from the outer leaflet of the plasma membrane, they get captured and disassembled by cell surface heparan sulfate proteoglycans resulting in the appearance of monomeric species FGF2 on cell surfaces (13–15). This process is based on the ability of heparan sulfates to compete against PI(4,5)P_2_ with an about hundred fold higher affinity for FGF2 compared with PI(4,5)P_2_ (14). The proposed mechanism has recently been confirmed in a fully reconstituted system using giant unilamellar vesicles (14) and is consistent with earlier observations demonstrating that membrane translocation depends on a fully folded state of FGF2 that permits PI(4,5)P_2_-dependent FGF2 oligomerization and interactions with heparan sulfate chains (27, 28). Recently, PI(4,5)P_2_- and heparan-sulfate-dependent translocation of FGF2 across the plasma membrane has also been visualized in living cells using single molecule TIRF microscopy. These studies revealed the real-time kinetics of this process with an average time interval for FGF2 membrane translocation of about 200 ms (6, 15).

In previous studies, we observed physical interactions between FGF2 and PI(4,5)P_2_ to be most efficient when PI(4,5)P_2_ was reconstituted in a lipid environment resembling plasma membranes. In particular, reconstituting PI(4,5)P_2_ in a pure PC lipid background or removing cholesterol from a plasma-membrane-like lipid composition resulted in decreased binding efficiency of FGF2 to PI(4,5)P_2_ (23, 24). In the light of cholesterol being known to exert profound effects on the organization of the plasma membrane with the lateral segregation into liquid-ordered and liquid-disordered domains being one example (29–33), we addressed a potential role of cholesterol in PI(4,5)P_2_-dependent FGF2 recruitment and translocation across the plasma membrane, the core step of its unconventional mechanism of secretion. We demonstrate in both biochemical reconstitution experiments and single molecule analyses in living cells that cholesterol enhances both PI(4,5)P_2_-dependent FGF2 recruitment at the inner plasma membrane leaflet and translocation into the extracellular space. Based on extensive atomistic molecular dynamics simulations, we find cholesterol to increase head group visibility of PI(4,5)P_2_ by exposing negative charges on the membrane surface in a way that promotes faster binding kinetics and a more stable interaction between FGF2 and PI(4,5)P_2_. Furthermore, we find cholesterol to induce clustering of PI(4,5)P_2_ molecules with the predominant appearance of trimers and tetramers. In a cellular context, at the inner plasma membrane leaflet, this phenomenon is likely to generate increased avidity enhancing PI(4,5)P_2_-dependent FGF2 oligomerization and membrane translocation to the cell surface. Finally, using droplet interface bilayers (DIB) inside a microfluidic setup, we measured bilayer tension as a function of the cholesterol concentration revealing a correlation between this parameter and the efficiency of PI(4,5)P_2_-dependent FGF2 membrane recruitment. Since an increase in bilayer tension is known to facilitate the formation of lipidic membrane pores (34–37), cholesterol may have a positive impact on PI(4,5)P_2_-dependent FGF2 membrane translocation in cells facilitating FGF2 oligomerization concomitant with the formation of toroidal membrane pores within the plasma membrane.

In conclusion, along with FGF2 interactions with the Na,K-ATPase (18, 19) and Tec kinase (20, 21) at the inner leaflet, our findings provide a compelling explanation for the high selectivity by which FGF2 is targeted to the plasma membrane, the subcellular site of FGF2 membrane translocation into the extracellular space.

## Results

### Cholesterol enhances PI(4,5)P_2_-dependent binding of FGF2 to lipid bilayers

In a previous study, we found that lipid bilayers made from a complex plasma-membrane-like lipid composition containing 2 mol% PI(4,5)P_2_ recruit FGF2 more efficiently than liposomes merely consisting of phosphatidylcholine (PC) and 2 mol% PI(4,5)P_2_ (23). To test a potential role for cholesterol in positively modulating PI(4,5)P_2_-dependent recruitment to lipid bilayers in a fully reconstituted system, we used a protein-lipid interaction assay based on analytical flow cytometry (24). To quantify FGF2 binding to PI(4,5)P_2_ in the context of increasing concentrations of cholesterol, we made use of a FGF2-Halo fusion protein labeled with Alexa Fluor 488 (AF488) and normalized binding efficiency by labeling liposomes with a rhodamine-coupled derivative of phosphatidylethanolamine (PE) (23, 24). In a first set of experiments, we analyzed the binding kinetics of FGF2-Halo-AF488 to liposomes containing 5 mol% PI(4,5)P_2_, 30 mol% cholesterol and 65 mol% PC (Fig. 1A). Using FGF2 tagged with a Halo domain, binding kinetics to PI(4,5)P_2_ were characterized by a linear phase until about 6 hours and a plateau being reached after about 12 hours of incubation. Based on these findings, we tested the impact of increasing concentrations of cholesterol in PI(4,5)P_2_-dependent FGF2 recruitment to membranes at different time points of incubation ranging from 1 hour to 24 hours (Fig 1B, subpanels a-d). These experiments revealed that increasing concentrations of cholesterol ranging from 10 to 50 mol% significantly enhance PI(4,5)P_2_-dependent FGF2 binding to membranes. This was particularly evident for incubation times of 1 and 3 hours (Fig. 1B, subpanels a and b) but was also detectable at 6 hours of incubation (Fig. 1B, subpanel c), a time span through which linear binding behavior was observed in the experiments shown in Fig. 1A. By contrast, following 24 hours of incubation at which PI(4,5)P_2_-dependent FGF2 binding was found to have reached a plateau (Fig. 1A), an impact of increasing concentrations of cholesterol could no longer be observed (Fig. 1B, subpanel d). Using a fully reconstituted system with all components purified to homogeneity, these findings demonstrate that cholesterol significantly enhances PI(4,5)P_2_-dependent FGF2 recruitment to lipid bilayers.

**Fig. 1:**
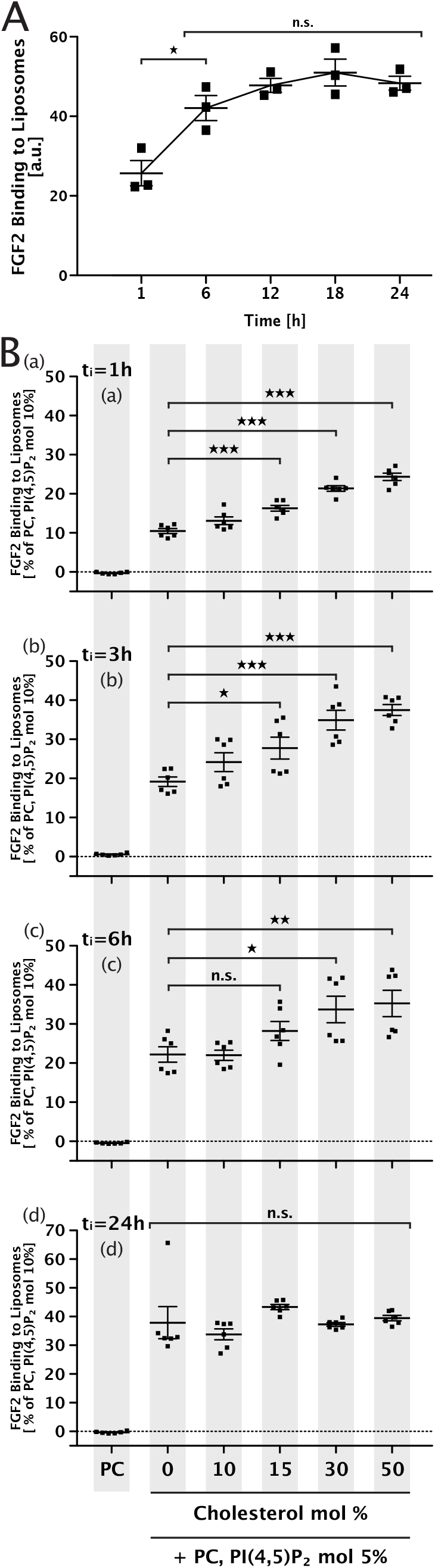
Cholesterol enhances PI(4,5)P_2_-dependent binding of FGF2 to lipid bilayers. FGF2-Halo-AF488 binding to PI(4,5)P_2_-containing liposomes was quantified using an analytical flow cytometry assay described previously (23, 24). Panel **A** displays a kinetic analysis using liposomes containing 5 mol% PI(4,5)P_2_, 30 mol% cholesterol and 65 mol% PC (PC5-CHOL30). Measurements were taken after 1, 6, 12, 18 and 24 hours of incubation. Panel **B** shows the quantitative analysis of FGF2-Halo-AF488 binding to various kinds of liposomes containing different levels of cholesterol after 1 (sub-panel a), 3 (sub-panel b), 6 (sub-panel c), and 24 hours (sub-panel d) of incubation. FGF2-Halo-AF488 binding to liposomes containing 10 mol% PI(4,5)P_2_ and 90 mol% PC (PC10 system; positive control) and liposomes consisting of 100 mol% PC (PC0 system; negative control) were used to normalize data. All data were corrected for background defined by binding of Halo-AF488 to the liposomal systems indicated. Standard errors (n=3 for A, n = 6 for B) and p-values are shown with ^n.s^p > 0.05; *p ≤ 0.05; ***p ≤ 0.001. The statistical analysis was based on a one-way ANOVA test.

### Cholesterol enhances head group visibility and clustering of PI(4,5)P_2_ on membrane surfaces

The electrostatic interaction of the headgroup of PI(4,5)P_2_ with the defined high affinity binding pocket in FGF2 is understood in great detail (14, 23). Using fully atomistic molecular dynamics simulations and free energy calculations, we aimed at revealing the mechanism by which cholesterol modulates FGF2 recruitment to PI(4,5)P_2_-containing lipid bilayers. Using extensive umbrella sampling simulations (38, 39), we quantified the free energy barriers associated with binding of FGF2 to POPC-based membranes containing 5 mol% PI(4,5)P_2_ in the presence and absence of 30 mol% of cholesterol. To avoid any bias due to different initial configurations, the systems were constructed by inserting a pre-formed FGF2-PI(4,5)P_2_ (1:4) complex in a bilayer with either 0 or 30 mol% of cholesterol (see Material and Methods for details). The two systems were first simulated for one microsecond at 298 K under NpT conditions using the GROMACS-2020 software (40) and the CHARMM36m forcefield (41). Thirty-nine windows with 0.1 nm spacing were constructed by gently pulling the protein away from the membrane in the z-direction (along the membrane normal direction) at 0.1 nm/ns and a force constant of 100 kJ mol^−1^ nm^−2^. To minimize membrane deformation during the pulling simulations, the head groups and tails of all membrane lipids were restrained on the z-axis using harmonic position restraints with a force constant of 10000 kJ mol^−1^ nm^−2^. The free energy profiles were analyzed as a function of the z distance (along the membrane normal direction) to the membrane surface (Fig. 2A), demonstrating the interaction between FGF2 and PI(4,5)P_2_ to represent a spontaneous process. This was true in both the presence and absence of cholesterol. However, cholesterol was found to exert a higher dissociation energy barrier shifting the energy minimum from −29 k_B_T to −35 k_B_T. Thus, in the presence of cholesterol, FGF2 binding to PI(4,5)P_2_ is both faster and more stable. These findings are consistent with the biochemical reconstitution experiments shown in Fig. 1.

**Fig. 2:**
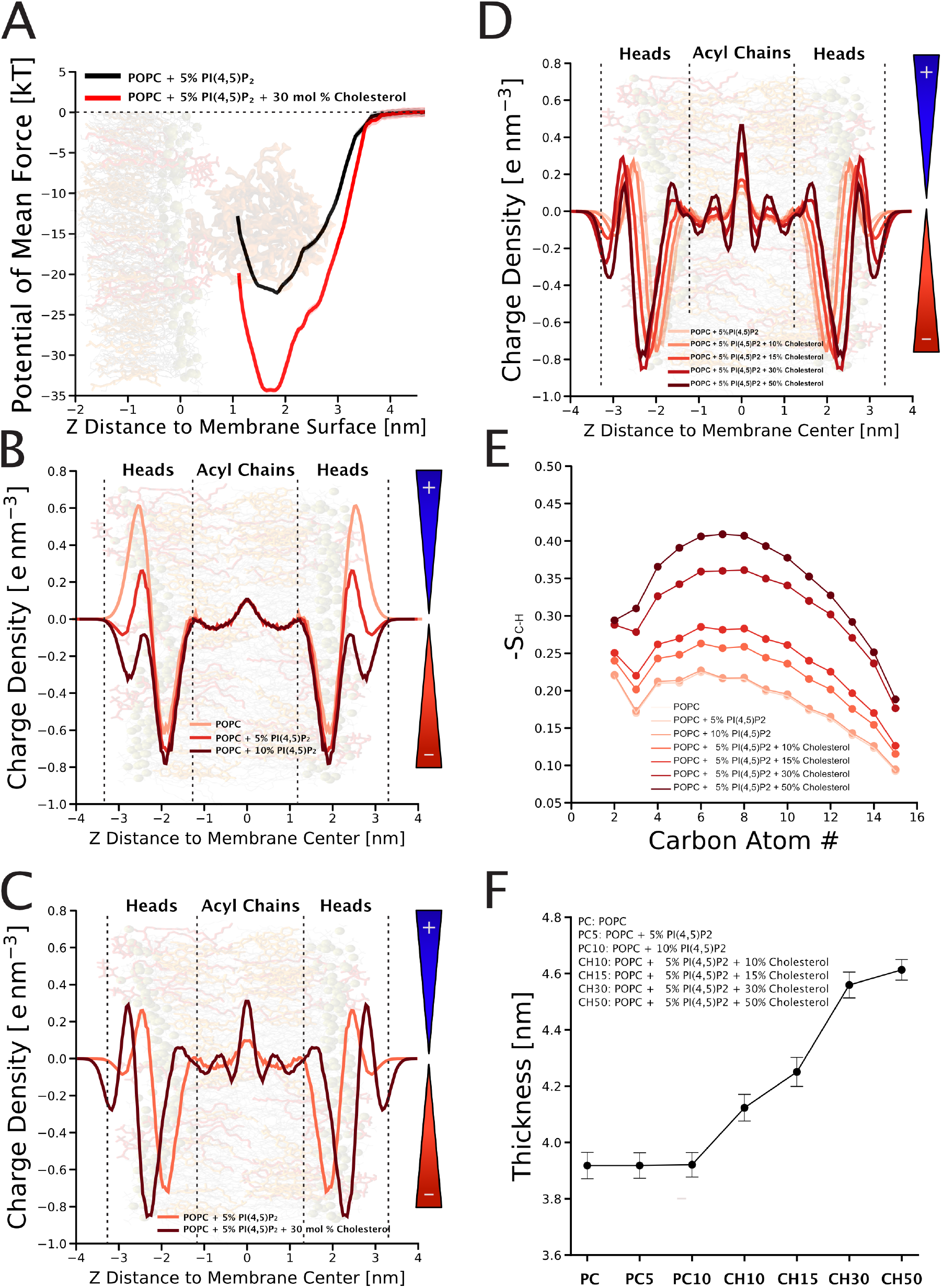
Cholesterol enhances PI(4,5)P_2_ head group visibility stabilizing FGF2 membrane recruitment. Panel **A** shows the potential of mean force (PMF) along with the z component of the distance between the center of masses of the protein and the P atoms of phospholipids in the closest leaflet. 39 windows with 0.1 nm spacing were simulated for 400 ns. The first 200 ns have been discarded from the PMF calculation and considered as equilibration time. Panels **B** and **C** show charge density profiles along the perpendicular axis of the bilayers, averaged over the last 500 ns of the pure bilayer simulations (no FGF2) without (panel **B**) and with 30 mol% of cholesterol (panel **C**). Panel **D** shows the charge density profiles for all lipid composition analyzed in this study. Panel **E** shows the deuterium order parameter (−S_CH_) of POPC sn-1 chains calculated from MD simulations for pure POPC membrane and for PI(4,5)P_2_-enriched (5 mol%) POPC-based membranes containing either 0, 10, 15, 30 or 50 mol% of cholesterol, respectively. S_CH_ was evaluated as averaged over the last 500 ns of simulations by taking both leaflets into account. Panel **F** shows the membrane thickness for a pure POPC membrane and PI(4,5)P_2_-enriched (5 mol%) POPC-based membranes containing either 0, 10, 15, 30 or 50 mol% of cholesterol, respectively. The thickness was calculated by measuring the z-component of the center of the mass distance between the two leaflets’ phosphate atoms. The data were averaged over the last 500 ns of the simulations with the error given as standard deviation.

To reveal the mechanism underlying the positive modulation of FGF2 binding to PI(4,5)P_2_ by cholesterol, we investigated the effect of cholesterol on structural and electrostatic properties of PI(4,5)P_2_-containing lipid bilayers (Fig. 2B and 2C). Seven different lipid compositions mimicking the *in vitro* experiments shown in Fig. 1 were analyzed (see Table 4 in Materials and Methods for simulations details). Intriguingly, we found that cholesterol affects the charge density distribution of lipid bilayers containing PI(4,5)P_2_ (Fig. 2A and 2B). These were calculated by summing up all charges per slice along the surface of lipid bilayers with the lipid compositions indicated (Fig. 2A and 2B). In a pure POPC bilayer (Fig. 2B, light-red line), the charge density shows two peaks in the headgroup region. The positive peak at +0,6 e/nm^3^ originates from the positive charge of the choline headgroup of POPC. The negative peak at −0,6 e/nm^3^ represents the negative net charges of phosphate groups. The substitution of POPC molecules with either 5 or 10 mol% of PI(4,5)P_2_ (Fig. 2B; red and dark-red lines) did not have any impact on the average charge density distribution in the phosphate region of membrane lipids. By contrast, a decrease of the positive charge density distributions in the region of the choline headgroups was observed from about +0.6 e/nm^3^ in the absence of PI(4,5)P_2_ to about +0.2 e/nm^3^ in the presence of 5 mol% of PI(4,5)P_2_ (Fig. 2B; red line) and about −0.3 e/nm^3^ for 10 mol% of PI(4,5)P_2_ (Fig. 2B; dark-red line).

In further simulations, we studied the effects of substituting POPC with cholesterol using lipid membranes containing 5 mol% PI(4,5)P_2_ (Fig. 2C and Fig. 2, supplement 1). Under these conditions, despite the relative content of PI(4,5)P_2_ remained the same, the average charge density in the headgroup regions became more negative with a maximum of about −0.3 e/nm^3^. Furthermore, at 30 mol% cholesterol, we observed the formation of transient PI(4,5)P_2_ clusters with trimers and tetramers being formed at the expense of monomers and dimers (Fig. 3 A and supplemental video 1). This phenomenon is documented in Fig. 3B in which the aggregation state of PI(4,5)P_2_ molecules is shown as a function of time. In this way, highly negatively charged spots of PI(4,5)P_2_ clusters are forming on the membrane surface that we propose to function in stabilizing FGF2 binding through increased avidity. Furthermore, when FGF2 binding to model membranes was simulated, 5 to 6 molecules of PI(4,5)P_2_ were observed to associate with FGF2 (Fig. 3, supplement 1 A-B and supplemental video 2), causing a strong local accumulation of negative charges that is likely to destabilize the lipid bilayer. This idea is supported by previous studies that have shown high charge-concentration gradients and electric fields across a membrane to induce pore formation (42, 43). The thermodynamic aspects of this phenomenon are likely to play a major role in the conversion of the lipid bilayer structure into a lipidic membrane pore during FGF2 oligomerization and membrane translocation.

**Fig. 3:**
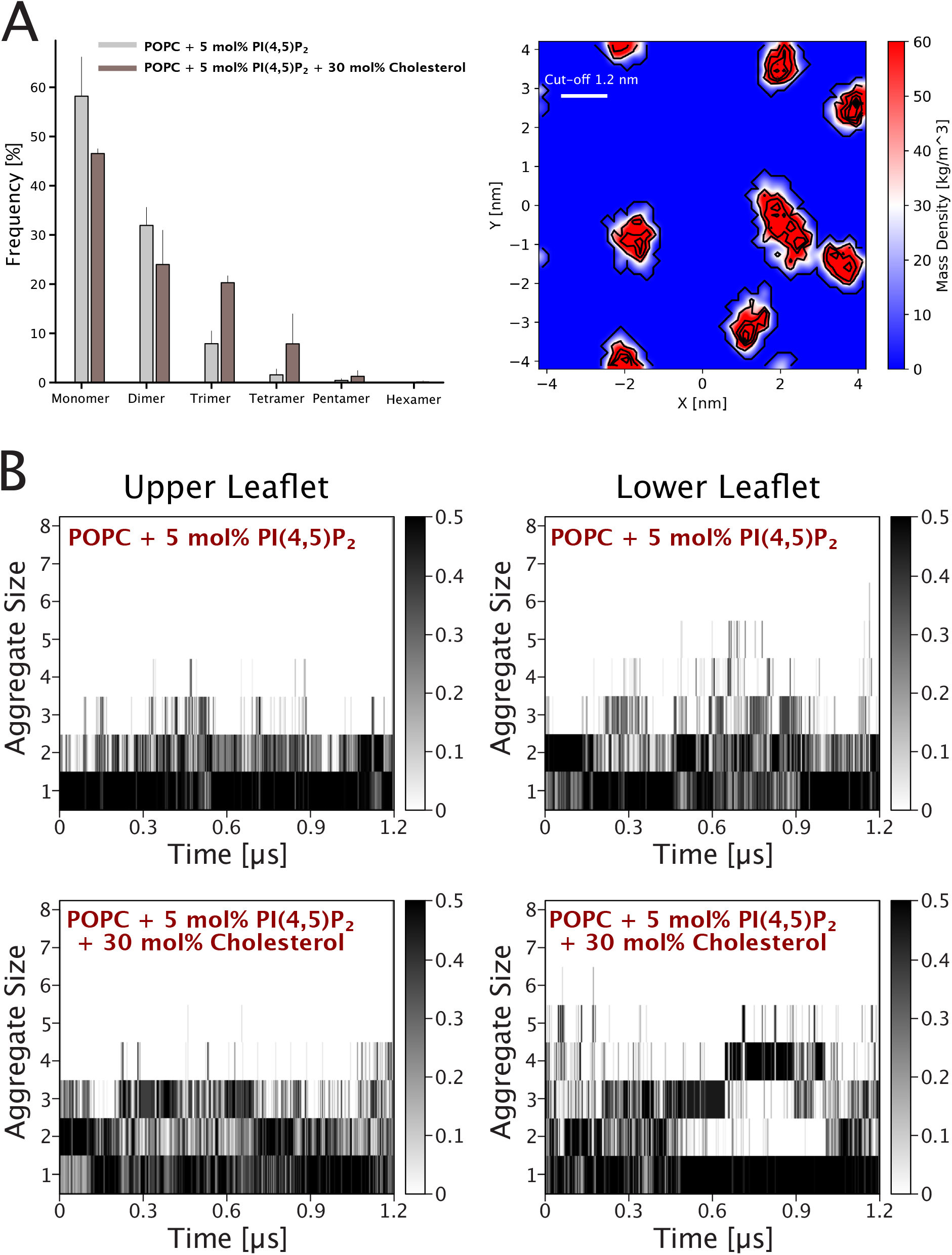
Cholesterol triggers clustering of PI(4,5)P_2_ molecules. Panel **A** shows a size-aggregation frequency analysis (right panel) for systems without (gray color) and 30 mol% of cholesterol (brown color). The analysis was averaged over the two membrane leaflets considering the last 1000 ns of simulations. Data are shown as the average of both bilayer leaflets, and the error is given as standard deviation. Two monomers were considered as a dimer if any of their atoms had a distance of less than 1.2 nm. The right panel represents a snapshot of a lateral partial density analysis with a dimer present as an example. Panel **B** provides the time evolution of the cluster fractional occupation number distribution of PI(4,5)P_2_ head groups for systems without (upper panels) and 30 mol% of cholesterol (bottom panels). The calculation was made for each membrane leaflet separately. Each panel represents a 1200 ns MD simulation of a system with 8 PI(4,5)P_2_ molecules (head groups only). The fractional occupation number was calculated every nanosecond and it is represented with a black and white scale bar.

### Cholesterol increases lateral bilayer tension in PI(4,5)P_2_-containing membranes

Combining microfluidic technology with drop shape analysis, we aimed at determining the role of cholesterol on membrane mechanics and its effect on bilayer tension (i.e. lateral membrane pressure). The presence of cholesterol is often associated with an increase in membrane tension, however, this phenomenon strongly depends on the specific membrane lipid species being analyzed (44–46). In the context of this study, we analyzed potential effects of cholesterol in the context of polydisperse lipid mixtures containing PI(4,5)P_2_, a non-bilayer lipid known to increase lateral pressure in the acyl chain regions of lipid bilayers (47). To determine this parameter, the bilayer contact angle θ and the interfacial tension were measured for each of the lipid compositions. To obtain θ, a lipid bilayer was formed at the intersection of a microfluidic set-up using a two-phase consisting of an aqueous buffer and squalene as a solvent for membrane lipids. Various lipid mixtures containing POPC, 5 mol% PI(4,5)P_2_, and cholesterol at either 0, 10, 15, 30 or 50 mol% were used to determine membrane tension. As shown in Fig. 4 – Supplement 1 A and Table 1, the bilayer contact angle θ was determined from optical micrographs as the angle between the two leaflets. For interfacial tension (σ) (Fig. 4 A-B), pendant buffer droplets in a squalene-lipid solution (1 mg/mL) were used as described in Materials and Methods.

**Table 1.** Values of bilayer contact angle (θ), interfacial tension (σ), bilayer tension (ϒ) and adhesion energy (E) for 5 lipid mixtures with different concentrations of cholesterol. The provides errors are standard deviations calculated using error propagation.

| Lipid ratio | $\theta$ (°) | $\sigma$ (mN/m) | $\gamma$ (mN/m) | $\epsilon$ (mJ/m<sup>2</sup>) |
| --- | --- | --- | --- | --- |
| POPC + 5% PI(4,5)P<sub>2</sub> | 31.0 ± 4.2 | 2.2 ± 0.2 | 3.8 ± 0.20 | 0.6 ± 0.2 |
| POPC + 5% PI(4,5)P<sub>2</sub> + 10% Chol | 29.4 ± 1.9 | 5.3 ± 0.5 | 9.2 ± 1.5 | 1.4 ± 0.4 |
| POPC + 5% PI(4,5)P<sub>2</sub> + 15% Chol | 30.6 ± 3.0 | 6.8 ± 1.2 | 11.7 ± 1.7 | 1.9 ± 1.0 |
| POPC + 5% PI(4,5)P<sub>2</sub> + 30% Chol | 22.9 ± 2.3 | 7.3 ± 0.3 | 13.4 ± 0.5 | 1.2 ± 0.1 |
| POPC + 5% PI(4,5)P<sub>2</sub> + 50% Chol | 21.5 ± 2.3 | 7.1 ± 0.8 | 13.2 ± 2.0 | 1.0 ± 0.2 |

**Fig. 4:**
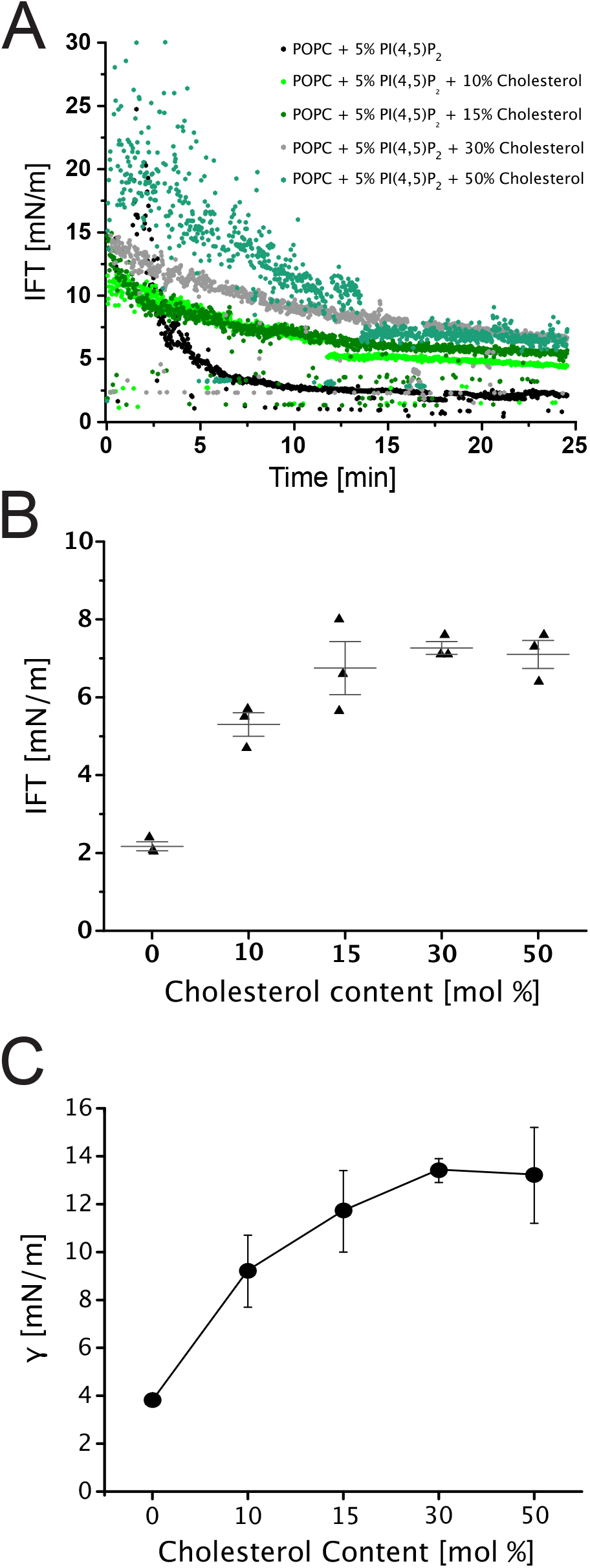
Cholesterol increases membrane tension in PI(4,5)P_2_-containing membranes. Panel **A** shows the interfacial tensions (ITFs) trends measured with pendant drop experiments as a function of time for POPC + 5% PI(4,5)P_2_, varying the cholesterol content (0, 10, 15, 30, 50 mol%). A buffer drop (HEPES 25 mM, KCl 150 mM) is immersed in a lipid-squalene solution (1 mg/mL) and its change in shape, that corresponds to the arrangement of the lipids at the newly created buffer-oil interface, is monitored. The IFT values recorded after reaching the plateau increase with higher cholesterol content. The IFT values given in panel **B** are average values (n=3) that were taken after reaching a plateau at t = 24 minutes. Errors are given as standard deviations. Panel **C** shows the values for bilayer tension ϒ of PI(4,5)P_2_-containing (5 mol%) membranes as a function of the cholesterol concentration. ϒ is calculated combining the bilayer angles obtained by analysis of the optical micrographs and the IFT values after the plateau through the equation 1. Errors were calculated using error propagation.

Using a microfluidic set-up (Fig. 4 – Supplement 1 B), we obtained a range of bilayer angles between 20° and 30° with a continuous decrease being observed as a function of rising concentrations of cholesterol (Fig. 4 – Supplement 1 A and Table 1).

An even more pronounced effect of cholesterol was observed for the interfacial tension values (σ) (Fig. 4B). In the absence of cholesterol, the value σ was found to be 2.2 ± 0.2 mN/m measured after reaching a plateau at t = 24 minutes (Fig. 4A). When cholesterol was added to the lipid mixtures, σ was found to increase to 5.3 ± 0.5, 6.8 ± 1.2, 7.3 ± 0.3, 7.1 ± 0.8 mN/m for cholesterol concentrations of 10, 15, 30 and 50 mol%, respectively. As shown in Fig. 4C, this trend is also reflected in the corresponding bilayer tension values (ϒ) combining σ and θ in equation 1 (see Materials and Methods). The highest bilayer tension value was found for lipid bilayers made from POPC, 5 mol% PI(4,5)P_2_ and 30 mol% cholesterol (ϒ=13.4 ± 0.5 Nm/m). Combining σ and θ in equation 2 (see Materials and Methods), the adhesion energies between the two monolayers were calculated (Table 1). When all lipid compositions were compared, a maximum value was observed for membranes containing 15 mol% cholesterol. At both lower and higher concentrations of cholesterol, the monolayer adhesion energy was found to be lower (Table 1). These observations are in line with the size of the contact areas found for different concentrations of cholesterol (Fig. 4 – Supplement 1 A and Table 1). Our findings establish a correlation between membrane tension and the cholesterol content for the lipid compositions used in the biochemical reconstitution experiments shown in Fig.1. Since an increase in tension is known to facilitate the formation of lipidic membrane pores (34–37), we propose cholesterol to increase membrane packing and tension in PI(4,5)P_2_-containing membranes, fostering FGF2 recruitment and translocation.

### Loading of cells with cholesterol enhances both FGF2 recruitment at the inner plasma membrane leaflet and FGF2 membrane translocation into the extracellular space

Using a single particle TIRF microscopy approach established previously (15, 18), we aimed at quantifying both FGF2 recruitment at the inner plasma membrane leaflet and FGF2 translocation to the cell surface under conditions of increased cholesterol levels in living cells (Fig. 5). This was achieved by using Cholesterol:Methyl-β-Cyclodextrin complexes to load cells with exogenous cholesterol (48, 49). We employed both CHO K1 and U2OS cells and analyzed cellular cholesterol levels by means of filipin staining and mass spectrometry (Fig. 5, supplement 1). Based on filipin staining, following treatment of cells with Cholesterol:Methyl-β-Cyclodextrin complexes, a significant increase of cholesterol levels by 56% in CHO K1 cells (Fig. 5, supplement 1, panels A and B) and 48% in U2OS cells (Fig. 5, supplement 1, panels D and E) could be observed. These findings were validated by mass spectrometry, demonstrating that treatment of cells with Cholesterol:Methyl-β-Cyclodextrin complexes resulted in increased cholesterol levels by 110% in CHO K1 cells and 97% in U2OS cells, respectively (Fig. 5, supplement 1, panels C and F). Thus, using two independent analytical methods, the described procedure was validated to increase cellular cholesterol levels in intact cells in a significant manner.

**Fig. 5:**
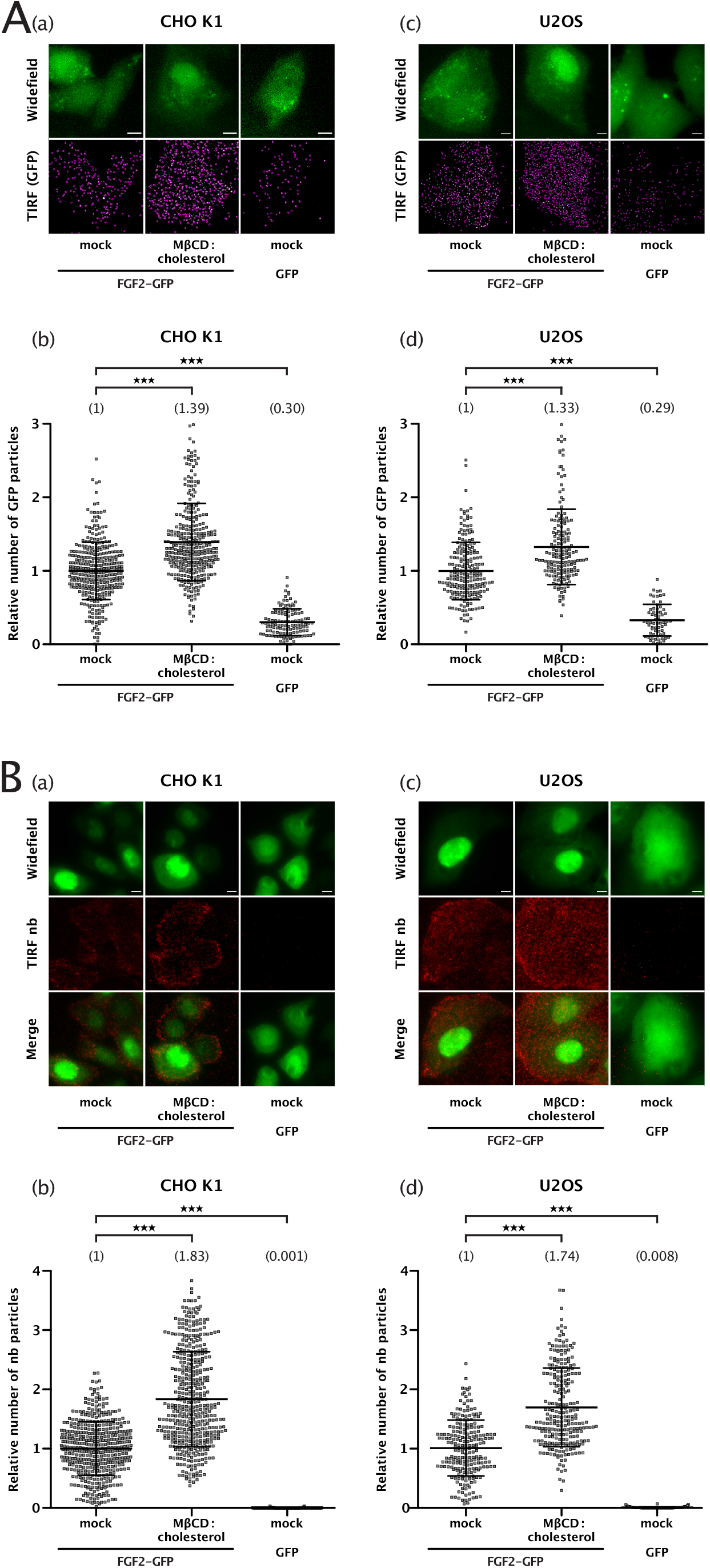
Increased cellular levels of cholesterol positively modulate FGF2 recruitment at the inner plasma membrane leaflet in living cells, as well as PI(4,5)P_2_-dependent FGF2 translocation to cell surfaces. CHO K1 and U2OS cell lines stably expressing either FGF2-GFP or GFP in a doxycycline-dependent manner were imaged by single molecule TIRF microscopy as described previously (15, 18). Before imaging, cells were treated with Cholesterol:Methyl-β-Cyclodextrin (1:10 molar ratio) complexes for 1 hour in culture conditions. Single FGF2-GFP or GFP particles were identified at the inner plasma membrane leaflet (**A**) (labelled by pink circles in subpanels **a**, **c**). For each condition, a widefield image and the first frame of the corresponding TIRF video are shown (subpanels **a**, **c**; Scale bar = 6 μm). Quantification of FGF2-GFP membrane recruitment at the inner leaflet of CHO K1 and U2OS cells is shown in subpanels **b** and **d**, respectively. Time-lapse TIRF movies with a total of 80 frames (100 ms/frame) were analyzed using the Fiji plugin TrackMate (71). The number of GFP particles were normalized for both surface area and the relative expression levels of FGF2-GFP for each experimental condition. Mean values are shown in brackets, with the mock condition set to 1. Data are shown as mean ± SD (n = 4). The statistical analysis was based on a one-way ANOVA test (***p ≤ 0.001). CHO K1 and U2OS cells were induced with doxycycline for 24h to express either FGF2-GFP or GFP (**B**). Following treatment with Cholesterol:Methyl-β-Cyclodextrin (1:10 molar ratio) complexes for 1 hour in culture conditions, cells were incubated on ice for 30 min with Alexa-Fluor-647-labeled anti-GFP nanobodies. After labelling of FGF2-GFP on cell surfaces, cells were fixed with 4% PFA at room temperature for 20 minutes and imaged by single molecule TIRF microscopy as established previously (15, 18). For each condition, a widefield image and the corresponding TIRF image is shown (subpanels **a**, **c**; Scale bar = 6 μm). Quantification of FGF2-GFP membrane translocation in CHO K1 and U2OS for all conditions indicated is shown in subpanel **b** and **d**, respectively. Single nanobody particles were analyzed using the Fiji plugin TrackMate (71). The number of nanobody particles detected per cell were normalized for the surface area of the corresponding cell. The mean values for each condition are shown in brackets, with the mock condition set to 1. Data are shown as mean ± SD (n = 4). The statistical analysis was based on a one-way ANOVA test (***p ≤ 0.001).

Based on the described procedures and experimental conditions, we quantified FGF2-GFP recruitment at the inner plasma membrane leaflet in intact CHO K1 and U2OS cells (Fig. 5A). For all conditions, both widefield and TIRF images were taken (Fig. 5A; subpanels a and c). While the widefield images allowed for the analysis of total expression levels of FGF2-GFP, the TIRF images were processed for the quantification of individual FGF2-GFP particles per surface area in the direct vicinity of the plasma membrane. These experiments demonstrated that treating cells with Cholesterol:Methyl-β-Cyclodextrin complexes to increase cellular cholesterol levels (Fig. 5, supplement 1) results in enhanced FGF2-GFP recruitment at the inner plasma membrane leaflet. This phenomenon could be observed in both cell lines being investigated with a significant increase of 39% in CHO K1 cells and 33% in U2OS cells, respectively (Fig. 5A, subpanels b and d). Similarly, under conditions of increased cellular cholesterol levels, we found FGF2-GFP translocation across the plasma membrane to be enhanced in a highly significant manner by 83% (CHO K1) and 74% (U2OS), respectively (Fig. 5B, subpanels b and d). These findings in intact cells are consistent with the biochemical reconstitution experiments shown in Fig. 1, demonstrating that cholesterol positively modulates FGF2 membrane recruitment by PI(4,5)P_2_, a process that triggers FGF2 translocation across the plasma membrane during unconventional secretion.

## Discussion

The current study originated from earlier observations suggesting the lipid environment to be an important modulator of PI(4,5)P_2_-dependent FGF2 recruitment to membrane surfaces (23, 24). Based on (i) biochemical reconstitution experiments, (ii) membrane tension analyses, (iii) extensive atomistic molecular dynamics simulations and (iv) cell-based FGF2 recruitment and secretion experiments using single molecule TIRF microscopy, we propose these effects to be linked to cholesterol, an abundant membrane lipid of plasma membranes.

We found PI(4,5)P_2_-dependent membrane recruitment of FGF2 to be affected by cholesterol in at least two ways. First, the presence of cholesterol has an impact on how negative charges of PI(4,5)P_2_ headgroups are presented on the membrane surface, promoting increased head group visibility for the PI(4,5)P_2_ binding site in FGF2. This effect is due to (i) exchange of PC molecules carrying a positively charged choline headgroup with cholesterol, mimicking the cholesterol gradient along the secretory pathway towards the plasma membrane (50–52) and (ii) the condensing effect of cholesterol on membrane surfaces that leads to an increased packing of lipids and a more dense arrangement of negative charges derived from PI(4,5)P_2_ (50, 53–57). In both molecular dynamics simulations and *in vitro* experiments presented in this study, cholesterol was added to membrane systems at the expense of POPC, a zwitter-ionic molecule which carries a positive charge in the headgroup surface region. Under these conditions, cholesterol affects membrane packing and tension, resulting in a phase transition from liquid-disordered to liquid-ordered arrangements as observed in molecular dynamics simulations (Fig. 2 E-F). The lateral segregation of membrane lipids with saturated and unsaturated fatty acids, respectively, and cholesterol into regions with high lipid packing (liquid-ordered domains) and less lipid packing (liquid-disordered domains) could be one principle that leads to a local enrichment of PI(4,5)P_2_ molecules (50, 52). Therefore, we propose that, in the presence of cholesterol, the negative charges of the headgroup of PI(4,5)P_2_ molecules might be packed more densely, facilitating electrostatic interactions with the PI(4,5)P_2_ binding pocket of FGF2. As demonstrated by free energy calculations, the described effects are linked to faster binding kinetics and a substantial stabilization of the interaction of FGF2 with PI(4,5)P_2_ with the free energy of binding increasing from −20 k_B_T to −35 k_B_T in the presence of cholesterol.

Second, we found cholesterol to induce clustering of PI(4,5)P_2_ molecules, a process that is likely to strengthen interactions between FGF2 and PI(4,5)P_2_ through increased avidity. In particular, in the presence of cholesterol, PI(4,5)P_2_ tends to form trimers and tetramers that we propose to enhance PI(4,5)P_2_-dependent FGF2 membrane recruitment. This process appears to be fostered by the ability of a single FGF2 molecule to bind five to six PI(4,5)P_2_ molecules through both the main high affinity binding site [defined by K127, R128 and K133; (14, 23, 26)] and additional low affinity PI(4,5)P_2_ binding sites on the positively charged surface of FGF2. Under these conditions, PI(4,5)P_2_-dependent oligomerization of FGF2 will result in a massive local accumulation of PI(4,5)P_2_ molecules at sites of FGF2 recruitment, a process that, due to the nature of PI(4,5)P_2_ as a non-bilayer lipid, is likely to exert membrane stress (47). This condition may facilitate FGF2 membrane translocation since, through the formation of a lipidic membrane pore with a toroidal architecture that allows for the accommodation of cone-shaped PI(4,5)P_2_ molecules (6, 8), this process is likely to relax bilayer stress at sites of PI(4,5)P_2_-dependent FGF2 recruitment and membrane translocation. This phenomenon might further be linked to variations of bilayer tension induced by cholesterol, a stress condition that is known to be relieved by the formation of a lipidic membrane pore (34–37). This is because the activation energy of tension-induced pore formation decreases with an increase in bilayer tension (58). Furthermore, the localized accumulation of negative charges derived from PI(4,5)P_2_ will increase the transmembrane electric field and, therefore, facilitate the emergence of lipidic membrane pores triggered by PI(4,5)P_2_-dependent FGF2 oligomerization (42, 43). In a cellular context, once membrane-spanning FGF2 oligomers have been inserted into the plasma membrane in a PI(4,5)P_2_-dependent manner, they will be captured and disassembled by cell surface heparan sulfates at the outer plasma membrane leaflet, thereby completing this unconventional secretory pathway with FGF2 being exposed on cell surfaces (5, 6, 13, 14).

In the specific context of FGF2 membrane recruitment and translocation, the results presented in this study offer a sound explanation for the high specificity by which the process of FGF2 membrane translocation into the extracellular space is physically linked to the plasma membrane, a subcellular structure that is enriched in both PI(4,5)P_2_ and cholesterol. Our findings are likely to be relevant to other cargo molecules of the UPS type I pathways of unconventional protein secretion that either directly or indirectly depend on PI(4,5)P_2_ (5–7, 59). This includes HIV-Tat and Tau, proteins that make direct physical contacts to PI(4,5)P_2_ as a prerequisite to enter their pathways of unconventional secretion (60, 61). In addition, under certain physiological conditions, unconventional secretion of IL-1β has been reported to follow a type I mechanism of UPS (62). Here, even though IL-1β does not bind to PI(4,5)P_2_ directly (63), membrane ruffles enriched in PI(4,5)P_2_ appear to play an important role in Gasdermin-dependent secretion of IL-1β (64), suggesting that so far unidentified interaction partners of PI(4,5)P_2_ play a role in unconventional secretion of IL-1β. Since the secretory mechanisms of HIV-Tat and Tau as well as Gasdermin-dependent secretion of IL-1β are all linked to the plasma membrane as the subcellular site of membrane translocation, the described role of cholesterol in positively modulating PI(4,5)P_2_-dependent FGF2 membrane recruitment is likely to extend to a larger group of UPS type I cargoes secreted by unconventional means. Furthermore, our findings are likely to have broad implications for the highly dynamic processes of phosphoinositide-dependent protein recruitment to membranes in general. This family of membrane lipids with differentially phosphorylated inositol head groups has been demonstrated to be distributed in a highly specific manner across different subcellular organelles (65). In this way, they have been implicated to exert functions in many cellular processes as diverse as endo- and exocytosis (66, 67), cell migration (68) and organellar contact sites (69), among others. Therefore, the highly dynamic interconversion of different types of phosphoinositides combined with different levels of cholesterol in various subcellular organelles might allow cells to fine-tune protein recruitment to membranes in a highly sophisticated manner to enable phosphoinositides to serve a broad variety of biological processes.

## Acknowledgements

This work was supported by grants from the Deutsche Forschungsgemeinschaft (WN; SFB/TRR 83, SFB/TRR 186 and DFG Ni 423/10-1, BB; SFB/TRR 186 and AG, JBF, HH, RF, KJ; SFB1027). AG thanks the Max Planck School “Matter of Life” supported by by the German Federal Ministry of Education and Research (BMBF) in collaboration with the Max Planck Society. IV thanks the Sigrid Juselius Foundation, the Academy of Finland (project no. 331349), Human Frontier Science Program (project no. RGP0059/2019), and the Helsinki Institute of Life Science (HiLIFE) Fellow program for financial support. FL and IV wish to acknowledge the CSC - IT Center for Science (Espoo, Finland) for computational resources. We would like to further acknowledge help from Holger Lorenz (ZMBH imaging facility) and Monika Langlotz (ZMBH FACS facility).

## Materials and Methods

### Recombinant proteins

A His-tagged FGF2-Halo fusion protein was expressed in *E. coli* BL21 Star cells and was purified to homogeneity in three steps using Ni-NTA affinity chromatography, heparin chromatography, and size exclusion chromatography (14). Desalting, buffer exchange and protein concentration were achieved by 30K ultra filtration (Amicon).

### Preparation of large unilamellar vesicles (LUVs)

All membrane lipids were purchased from Avanti Polar Lipids that were either purified from natural extracts [bovine liver (PC), porcine brain (PI(4,5)P_2_), and ovine wool (cholesterol)] or were made as a synthetic product (Rhodamine-labeled PE). Liposomes were prepared as described previously (22–24, 26). In brief, chloroform-dissolved lipid mixtures containing (750 μl containing 3 μmol total lipids) were dried in round bottom flasks at 25°C under vacuum to yield a homogeneous lipid film. To obtain a final lipid concentration of 4 mM, dried lipid films were slowly resuspended in 750 μl HK buffer (150 mM KCl and 25 mM HEPES, pH = 7.4) containing 10 % (w/v) sucrose. The obtained lipid mixtures were subjected to ten freeze/thaw cycles (50°C/liquid nitrogen) to produce a preparation of unilamellar liposomes. A size-distribution of 250–300 nm in diameter was obtained by 37 size-extrusion steps using a mini-extruder (Avanti Polar Lipids) equipped with a 400 nm filter. The final size distribution of liposomes was found to have an average diameter of 259 ± 32 nm as analyzed by dynamic light scattering (Wyatt). Different lipid compositions were used to prepare liposomes, all of which contained 1 mol% Rhodamine-PE (Table 1).

### Quantification of protein-lipid interactions using analytical flow cytometry

Using a FGF2-Halo fusion protein fluorescently labeled with AF488, protein binding to PI(4,5)P_2_-containing liposomes was analyzed by analytical flow cytometry (Becton-Dickinson FACS Calibur along with data analysis using CellQuest Pro software) as described previously (23, 24). Briefly, liposomes (final lipid concentration = 1 mM) were blocked with 4% (w/v) fatty-acid-free BSA (Roche) in HK buffer for one hour at 25°C. FGF2-Halo-AF488 (5 μM) was incubated at 25°C for the time indicated. Following extensive washing with HK buffer (150 mM KCl and 25 mM HEPES, pH = 7.4), FGF2-Halo-AF488 binding to PI(4,5)P_2_-containing lipid bilayers was recorded for 30.000 individual liposomes detected by rhodamine-PE fluorescence and light scattering. Primary data were corrected for background fluorescence defined by unspecific binding of AF488-labeled Halo (lacking FGF2) to liposomes. FGF2-Halo-AF488 binding signals derived from liposomes containing 10 mol% of PI(4,5)P_2_ and 90 mol% PC (PC10) were set to 100 % and used as a reference value. Signals derived from PC-only containing liposomes (PC0) were used as the negative control.

### Preparation of Cholesterol:Methyl-β-Cyclodextrin complexes

Chloroform-dissolved cholesterol was dried under a stream of nitrogen and, subsequentially, under vacuum at room temperature in glass tubes. A Methyl-β-Cyclodextrin solution (200 mM) in Live Cell Imaging Solution (Thermo Fisher Scientific) was added to the cholesterol lipid film at a molar ratio of 10 to 1. The mixture was sonicated and vortexed for one hour at room temperature, resulting in a complete dissolution of the cholesterol lipid film. Aliquots were stored at −20 °C. Cells were treated with Cholesterol:Methyl-β-Cyclodextrin complexes (final cholesterol concentration = 0.2 mM) for 1 hour at 37°C.

### Single-molecule TIRF microscopy

To quantify both FGF2-GFP recruitment at the inner plasma membrane leaflet and translocation to cell surfaces, a previously established single particle TIRF assay was employed (15, 18). Widefield fluorescence and TIRF images were acquired using an Olympus IX81 xCellence TIRF microscope equipped with an Olympus PLAPO×100/1.45 Oil DIC objective lens and a Hamamatsu ImagEM Enhanced (C9100-13) camera. Data were recorded and exported in Tagged Image File Format (TIFF) and analyzed via Fiji (70). GFP fluorescence was excited with an Olympus 488 nm, 100 mW diode laser.

For the quantification of FGF2-GFP recruitment at the inner leaflet of the plasma membrane, CHO K1 and U2OS cells were seeded in μ-Slide 8 Well Glass Bottom plates (ibidi) (15, 18). Before imaging was started, cells were incubated with Cholesterol:Methyl-β-Cyclodextrin complexes for 1 hour at 37°C. The quantification of FGF2-GFP particles recruitment to the plasma membrane was achieved through the analysis of time-lapse TIRF movies. The frame of each cell was selected by widefield imaging. The number of FGF2-GFP particles were normalized to the cell surface area (μm^2^) and to the expression level of FGF2-GFP. The latter was quantified for each individual cell at the first frame of each time-lapse TIRF movie using ImageJ. The total number of FGF2-GFP particles per cell was quantified employing the Fiji plugin TrackMate (71).

For the quantification of FGF2-GFP translocation to cell surfaces, CHO K1 and U2OS cells were seeded in μ-Slide 8 Well Glass Bottom plates (ibidi) followed by incubation for 24 h in the presence of 1 μg/ml doxycycline to induce FGF2-GFP expression. Prior to the addition of Cholesterol:Methyl-β-Cyclodextrin complexes, cells were washed with heparin (Sigma; 500 μg/ml) to remove FGF2-GFP particles from cell surfaces. Cells were then treated with Cholesterol:Methyl-β-Cyclodextrin complexes (final cholesterol concentration = 0.2 mM) for 1 hour at 37°C. Afterwards, the medium was removed and cells were rinsed twice with Live Cell Imaging Solution (Thermo Fisher Scientific). Cells were further incubated on ice with membrane impermeable Alexa Fluor 647– labeled anti-GFP nanobodies (Chromotek) for 30 min. Afterwards, they were rinsed three times with Live Cell Imaging Solution and fixed with 4% PFA (Electron Microscopy Sciences) for 20 min at room temperature. Nanobody fluorescence was excited with an Olympus 640 nm, 140 mW diode laser. The quantification of FGF2-GFP on cell surfaces was achieved through a quantitative analysis of TIRF images. The frame of each cell was selected by widefield imaging. The number of nanobody particles were normalized to the cell surface area (μm^2^). The total number of nanobody particles per cell was quantified employing the Fiji plugin TrackMate (71). Background fluorescence was subtracted for all representative images shown.

### Quantification of cellular cholesterol levels based on filipin imaging

Following treatment with Cholesterol:Methyl-β-Cyclodextrin complexes, cellular cholesterol levels were quantified by filipin (Sigma) staining and confocal imaging. CHO K1 and U2OS cells were seeded in μ-Slide 8 Well Glass Bottom plates (ibidi). After 24 hours of incubation, cells were incubated for 1 hour with Cholesterol:Methyl-β-Cyclodextrin complexes. Cells were then rinsed three times with PBS and fixed with 3% PFA for 1 hour at room temperature. Afterwards, to quench PFA, cells were rinsed three times with PBS and incubated with 1.5 mg/mL glycine in PBS for 10 min at room temperature. A stock solution of filipin (25 mg/mL in DMSO) was prepared and diluted to prepare a working solution of 50 μg/mL in PBS, supplemented with 10% FCS. Cells were stained with filipin for 2 hours at room temperature and rinsed three times with PBS. Imaging was performed using a confocal laser scanning microscope (Zeiss LSM800) with a 63× oil objective.

### Quantification of cellular cholesterol levels using mass spectrometry

Cells were subjected to lipid extractions using an acidic Bligh & Dyer (72). Lipid standards were added prior to extractions, using a master mix containing 50 pmol phosphatidylcholine (PC)-standard mix (PC 13:0/13:0, 14:0/14:0, 2.0:0/20:0; 21:0/21:0; Avanti Polar Lipids) and 150 pmol D_7_-cholesterol (Avanti Polar Lipids). The final CHCl_3_ phases were evaporated under a gentle stream of nitrogen. Lipid extracts were resuspended in 60 μl 10mM ammonium acetate in methanol and diluted 1:4 in 96-well plates (Eppendorf twin.tec^®^ 96; Sigma-Aldrich) prior to measurement of PC species, applying precursor ion scanning (prec) in positive ion mode (+prec184, mass range: m/z 644-880). For cholesterol determination, the remaining lipid extract was evaporated and subjected to acetylation as described^2^. Following acetylation, samples were evaporated and resuspended in 60 μl 10mM ammonium acetate in methanol, diluted 1:10 in 96-well plates (Eppendorf twin.tec^®^ 96; Sigma-Aldrich). Cholesterol measurements were performed in positive ion mode, applying neutral loss (NL) scanning (+NL 77, mass range: m/z 442-480, for MS parameters see Table 1) on a QTRAP 5500 (Sciex; Canada) mass spectrometer using chip-based (HD-D ESI Chip; Advion Biosciences; USA) electrospray infusion and ionization via a Triversa Nanomate (Advion Biosciences; USA) as previously described (73).

The scan rate was set to 200 Da/s with a step size of 0.1 Da. Data evaluation was done using LipidView 1.3 (Sciex; Canada) and a software developed in-house (ShinyLipids). One sample (Kon A) contained only the standards and was used for background subtraction. Amounts of endogenous lipid species were calculated on the basis of the intensities of internal standards. PC is a bulk lipid of cellular membranes and was used as reference for cholesterol determinations.

### Molecular Dynamics Simulations

Atomistic molecular dynamics (MD) simulations were performed using the CHARMM36m force field for lipids and proteins, the CHARMM TIP3P force field for water, and the standard CHARMM36 force field for ions (41). The GROMACS 2020 simulation package (40) was used in all simulations. For FGF2, we used structural information based on PDB ID: 1BFF that covers the crystal structure of residues 26 to 154 of the monomeric form of FGF2 (74). The N- and C-termini were modelled as charged residues. To match the experimental membrane systems used in Fig. 1, the CHARMM-GUI webserver (75) was used to model seven different lipid compositions (Table 2) with 300 lipids each. All membrane systems were first simulated for 1.2 microseconds. The final snapshot for each membrane lipid composition was used to build the protein-membrane system. FGF2 was placed two nanometers far from the membrane surface at ten different random orientations (see Table 2 for simulations details). The protein-membrane systems were first energy-minimized in vacuum and later hydrated and neutralized by an appropriate number of counter-ions, followed by the addition of 150 mM potassium chloride to mimic the experimental conditions. All systems were first energy-minimized and an equilibration step was used to keep the temperature, pressure, and number of particles constant (NpT ensemble). During this step, proteins were restrained in all dimensions, whereas the first heavy-atom of each lipid was restrained in the xy-plane of the membrane with a force constant of 1000 kJ/mol.

**Table 2.**
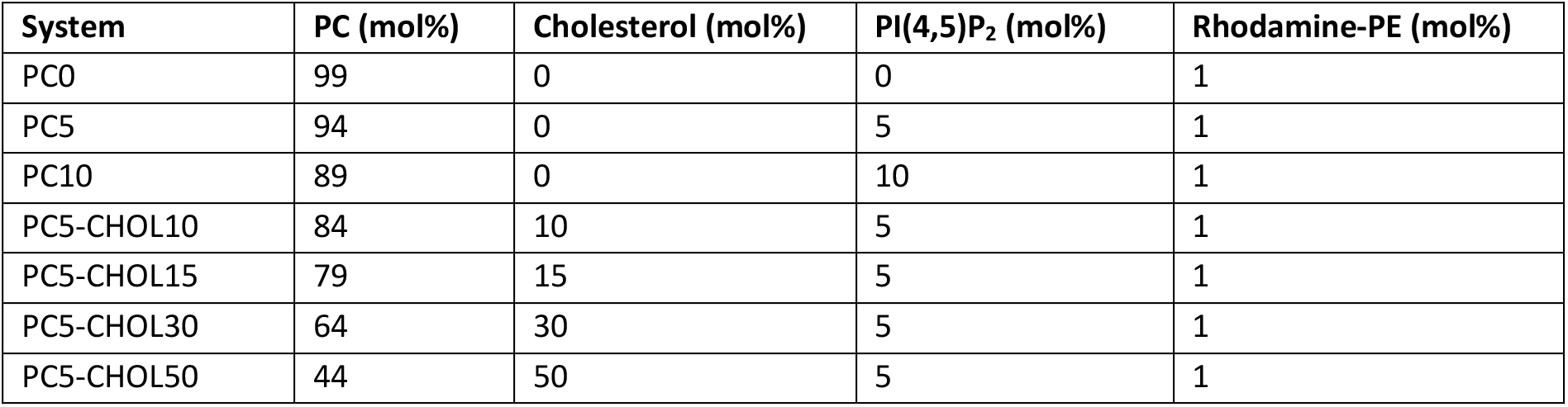
Liposome compositions used for preparing unilamellar liposomes

**Table 3:** MS parameters. CAD, collisionally-activated dissociation (N_2_); CE, collision energy; Chol, cholesterol; CUR, curtain gas (N_2_); CXP, collision cell exit potential; DP, declustering potential; EP, entrance potential; IHT, interface heater temperature; MCA, multi-channel acquisition; PC, phosphatidylcholine; Q, quadrupole.

| Lipid class | DP [eV] | EP [eV] | CE [eV] | CXP [eV] | CUR [psi] | IHT [°C] | CAD [psi] | MCA | Q1 | Q3 |
| --- | --- | --- | --- | --- | --- | --- | --- | --- | --- | --- |
| PC | 100 | 12 | 32 | 13 | 20 | 60 | 5 | 300 | high | unit |
| Chol | 100 | 10 | 12 | 14 | 20 | 60 | 5 | 600 | unit | unit |

**Table 4.** Details on the atomistic simulations. Lipid names correspond to the CHARMM-GUI lipid model. PI(4,5)P_2_ corresponds to SAPI25, Cholesterol to CHL1, and water to TIP3P.

| System | # POPC | # PI(4,5)P<sub>2</sub> | # Chol | # Water | # K<sup>+</sup> | # CL<sup>-</sup> | # Repeats | Time (ns) |
| --- | --- | --- | --- | --- | --- | --- | --- | --- |
| PC0 | 300 | 0 | 0 | 34575 | 94 | 106 | 10 | 1200 |
| PC5 | 284 | 16 | 0 | 34516 | 146 | 94 | 10 | 1200 |
| PC10 | 270 | 30 | 0 | 34770 | 203 | 95 | 10 | 1200 |
| PC5-CHOL10 | 254 | 16 | 30 | 33363 | 143 | 91 | 10 | 1200 |
| PC5-CHOL15 | 238 | 16 | 46 | 29281 | 132 | 80 | 10 | 1200 |
| PC5-CHOL30 | 194 | 16 | 90 | 24707 | 119 | 67 | 10 | 1200 |
| PC5-CHOL50 | 134 | 16 | 150 | 22619 | 114 | 62 | 10 | 1200 |

The Nose-Hoover thermostat (76) was used to maintain the temperature at 298 K with a time constant of 1.0 ps. The pressure of 1 atm was kept constant using the Parrinello–Rahman barostat (77) with a time constant set to 5.0 ps and isothermal compressibility to a value of 4.5 × 10^−5^ bar^−1^. The semi-isotropic pressure-coupling scheme was used. For neighbor searching, we used the Verlet scheme (78) with an update frequency of once every 20 steps. Electrostatic interactions were calculated using the Particle Mesh Ewald method (79). Periodic boundary conditions were applied in all directions. The simulations were carried out using an integration time step of 2 fs until 1200 ns were reached. The first 500 ns of all analyses were considered as equilibration time and excluded from the analysis. The total time scale of the simulations (including free energy calculations) was about 115 μs.

### Free Energy Calculations

The calculation of the potential of mean force (PMF) for the interaction between FGF2 and PI(4,5)P_2_ in POPC-based membranes was calculated using the umbrella sampling technique (38, 39). The systems were built by inserting, using Javanainen’s method (80), a pre-formed FGF2-PI(4,5)P_2_ (1:4) complex in two membranes with 0 and 30 mol% of cholesterol, respectively. FGF2-PI(4,5)P_2_ (1:4) complexes were built from an unbiased molecular dynamics simulation of FGF2 bound to PI(4,5)P_2_ in its high-affinity orientation (14), using a POPC-based bilayer containing 5 mol% of PI(4,5)P_2_. Before starting the pulling simulations, the systems were minimized and simulated for 500 ns to stabilize the FGF2-PI(4,5)P_2_ (1:4) complex bound to the lipid bilayer. Thirty-nine windows with 0.1 nm spacing were constructed by pulling FGF2 on the z-axis (along the membrane normal direction) with a pulling rate of 0.1 nm/ns and a force constant of 100 kJ mol-1 nm-2. All lipids were restrained in the xy-plane applying harmonic position restraints with a force constant of 10000 kJ mol^−1^ nm^−2^. Each window was simulated for 400 ns with the first 200 ns being considered an equilibration phase that was discarded from the free energy calculations. The PMF was reconstructed using the weighted histogram analysis method implemented in GROMACS (81). The reconstructed free energy profiles’ statistical errors were estimated with the Bayesian bootstrap method (82) using 200 bootstrap iterations.

### Analysis Tools

The charge density profiles shown in Fig. 2 and Fig.2-Supplement 1 were calculated with the gmx density GROMACS tool employing -symm, -center, and -d Z options. The box has been divided into 200 slices. The lateral density profile shown in Fig. 3A (right panel) was calculated with the g_mydensity tool (83). For the cluster analysis shown in Fig. 3A and 3B, we employed the g_aggregate tool (83) and in-house python script, using a minimum distance cut-off of 1.2 nm. The adaptive Poisson-Boltzmann Solver (APBS) calculation of the electrostatic surface of FGF2 shown in Fig 3-Supplement 1A was achieved using the APBS web-server (84). All images, snapshots, and movies were rendered with the Visual Molecular Dynamics (VMD) software (85).

### Microfluidics

To form a freestanding lipid membrane two microfluidic cross channel geometries previously tested as a platform to produce stable bilayers were employed (86–88). In both geometries, a continuous phase consisting of lipids dissolved in squalene solution (5 mg/mL) separates two fingers of HEPES buffer (25 mM, KCl 150 mM). Once buffer fingers are injected in the device with the lipid solution previously injected, the lipids will decorate the interface between the two phases forming two monolayers that are not in contact (Fig. 4 – Supplement 1 B). A slow flow is created by hydrostatic pressure such that the lipid decorated buffer fingers are brought in close vicinity until they make a direct contact. When the two monolayers face each other, a zipping phenomenon driven by the intermolecular forces between the hydrophobic chains will occur and the bilayer will form (87). In this setup, the control of pressure and lipid concentrations is necessary to avoid rupture of the lipid bilayer (89). The lipid bilayer contact angle θ was then obtained by analyzing optical micrographs. To guarantee the stability of the fingers, a good wettability of the phases was required. Therefore, a hydrophobic chip of PDMS was used and prepared as described previously (86, 90). Briefly, silicon elastomer base and curing agent (Sylgard 184, Dow Corning, USA) were mixed in a ratio of 10:1 and were casted on a silicon wafer, reproducing negative structures fabricated by photolithography with a SU-8 photoresist, and cured for 6 h at 65 °C. The prepared chip and a PDMS coated glass slide were then exposed to nitrogen plasma treatment (Diener electronic GmbH, Germany), sealed and heated at 125 °C for 1-2 h to restore hydrophobic properties. Optical images of bilayer formation occurring in the chip were acquired with SciDAVis or Zeiss software at 10x magnification (Zeiss observer Z1 and Z7 with pco camera Imager pro X).

### Interfacial tension measurements

Interfacial tension σ of a single lipid monolayer was measured via the standard pendant drop method using a OCA20 goniometer (OCA 20, DataPhysics Instruments GmbH, Filderstadt, Germany). Briefly, a 1 mg/mL solution of lipid in squalene was stirred at 45 °C for three hours and transferred into an optical glass chamber. A drop of HEPES buffer (25 mM, KCl 150 mM) was formed at the end of a hanging needle that was immersed in the lipid solution. The shape change of the pendant drop was recorded over time. To obtain the interfacial tension σ, its shape was fitted according to the Young-Laplace equation. The decrease of σ due to the adsorption of lipids to the newly created interface was recorded over 20-30 minutes until a plateau was reached.

### Bilayer Tension and Adhesion Energy

From the values for the interfacial tension σ and the bilayer contact angle θ, the bilayer tension ϒ was calculated using Young’s equation (91, 92):

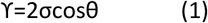

Furthermore, the corresponding adhesion energy ε per unit area between the bilayer sheets was calculated from the values for interfacial tension and bilayer contact angle using the Young−Dupré relation:

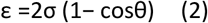

---

**Fig. 3 – Supplement 1:**
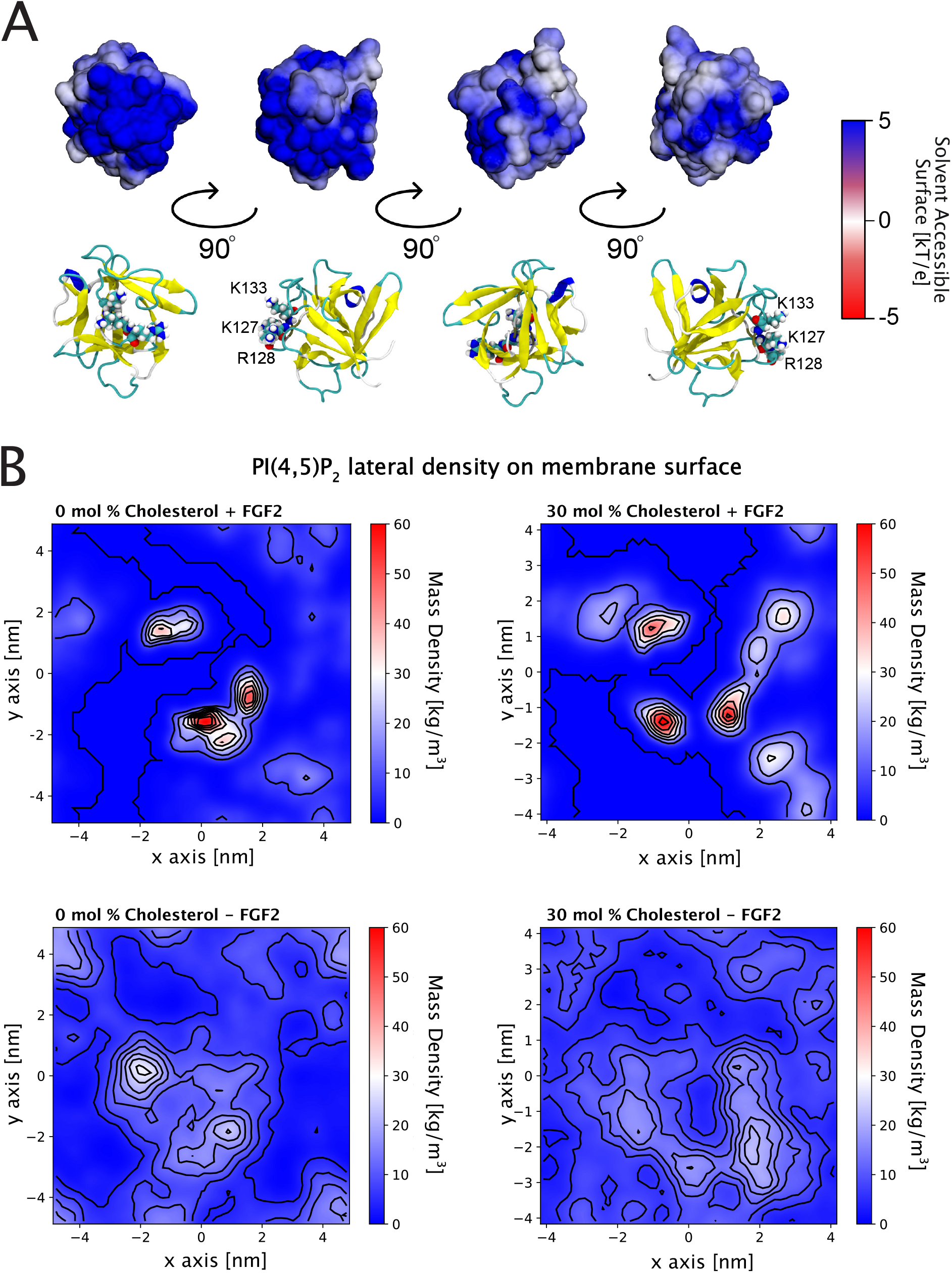
FGF2 binds to five to six PI(4,5)P_2_ molecules within different surface areas in a cholesterol-independent manner. Adaptive Poisson-Boltzmann Solver (APBS) calculations on the electrostatic surface of FGF2 (panel **A**) and localization of PI(4,5)P_2_ head groups expressed as mass density for the systems containing either 0 or 30 mol% cholesterol in the presence (upper panel) or absence (lower panel) of interactions with FGF2 (panel **B**). Data were averaged over the last 500 ns of the simulations.

**Fig. 3 – Supplemental Video 1:**

Time evolution of the lateral partial mass density analysis of PI(4,5)P_2_ head groups for membrane systems containing either 0 (left panel) or 30 mol% of cholesterol (right) in the absence of FGF2.

**Fig. 3 – Supplemental Video 2:**

Time evolution of the system with 30 mol% of cholesterol in presence of FGF2 (right panel) and its lateral partial mass density evolution of PI(4,5)P_2_ head groups.

**Fig. 4 Supplement 1:**
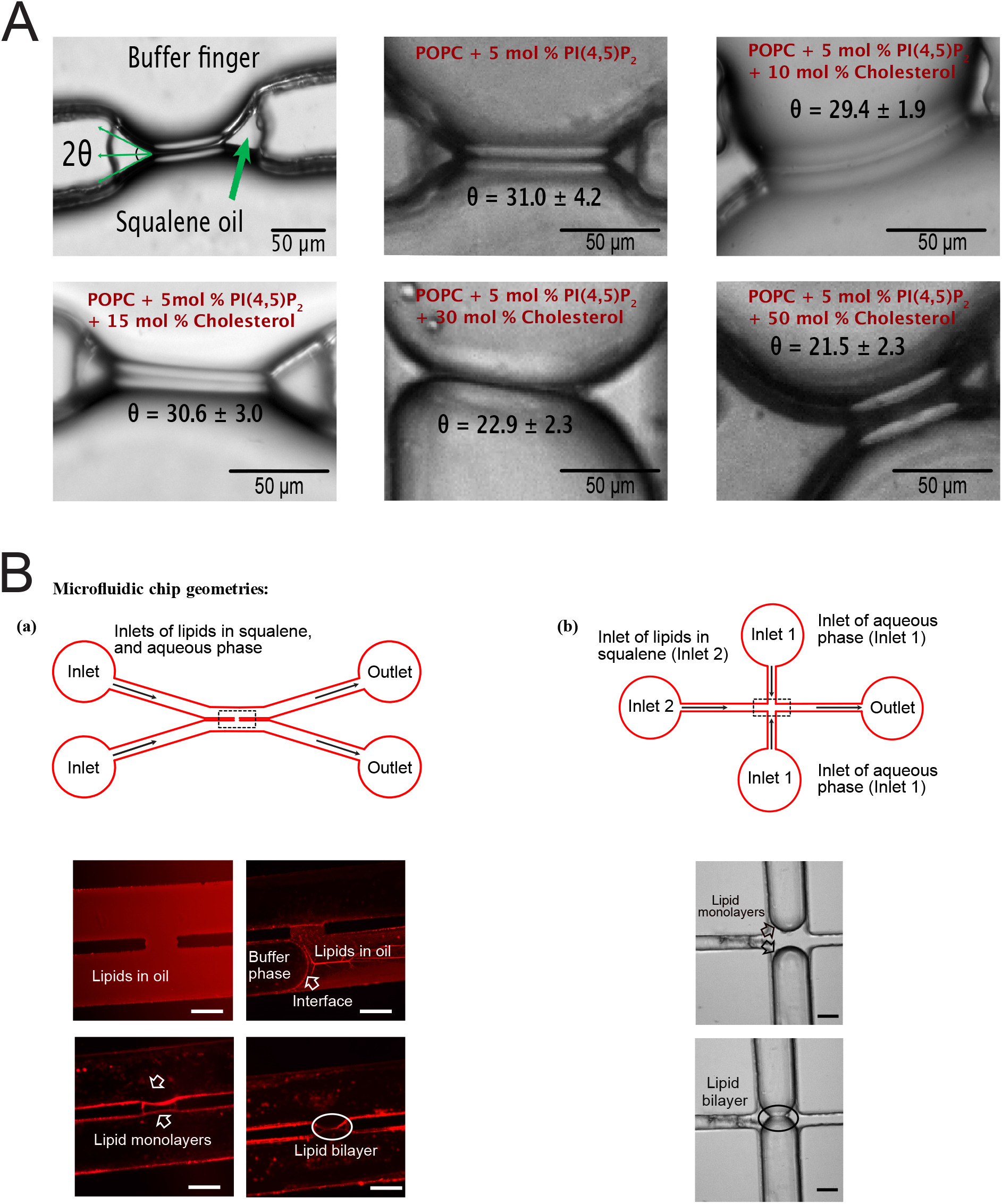
Analysis of bilayer contact angles using a microfluidics setup. Panel **A** shows optical micrographs of the bilayer contact angles for artificial membranes with different cholesterol contents increasing from top to bottom (scale bar = 50 μm). Panel **B** shows a schematic overview of the chip geometries used. Both the devices have a cross-geometry and the lipid monolayers meet at the intersection of the channels forming the bilayer. In subpanel **a**, fluorescence microscopy micrographs show how the chip was first filled by injection from one of the two inlets with a lipid solution in squalene stained with Rhodamine-labeled PE and then with a buffer phase using hydrostatic pressure. This setup allowed for the lipids to decorate the oil-buffer interface. In subpanel **b**, the chip was also filled with membrane lipids (inlet 2) as part of a squalene solution containing Rhodamine-labeled PE. Afterwards, the two buffer fingers were introduced from opposite sides (inlets 1) resulting in membrane lipid decoration of the two interfaces. After generating a close contact through hydrostatic pressure manipulation, the membrane lipids formed a bilayer as shown in the optical micrographs. The geometry of the chip in subpanel **a** was found to form more stable bilayers due to a better control of hydrostatic pressure. The scale bar is 75 μm.

**Fig. 5 – Supplement 1:**
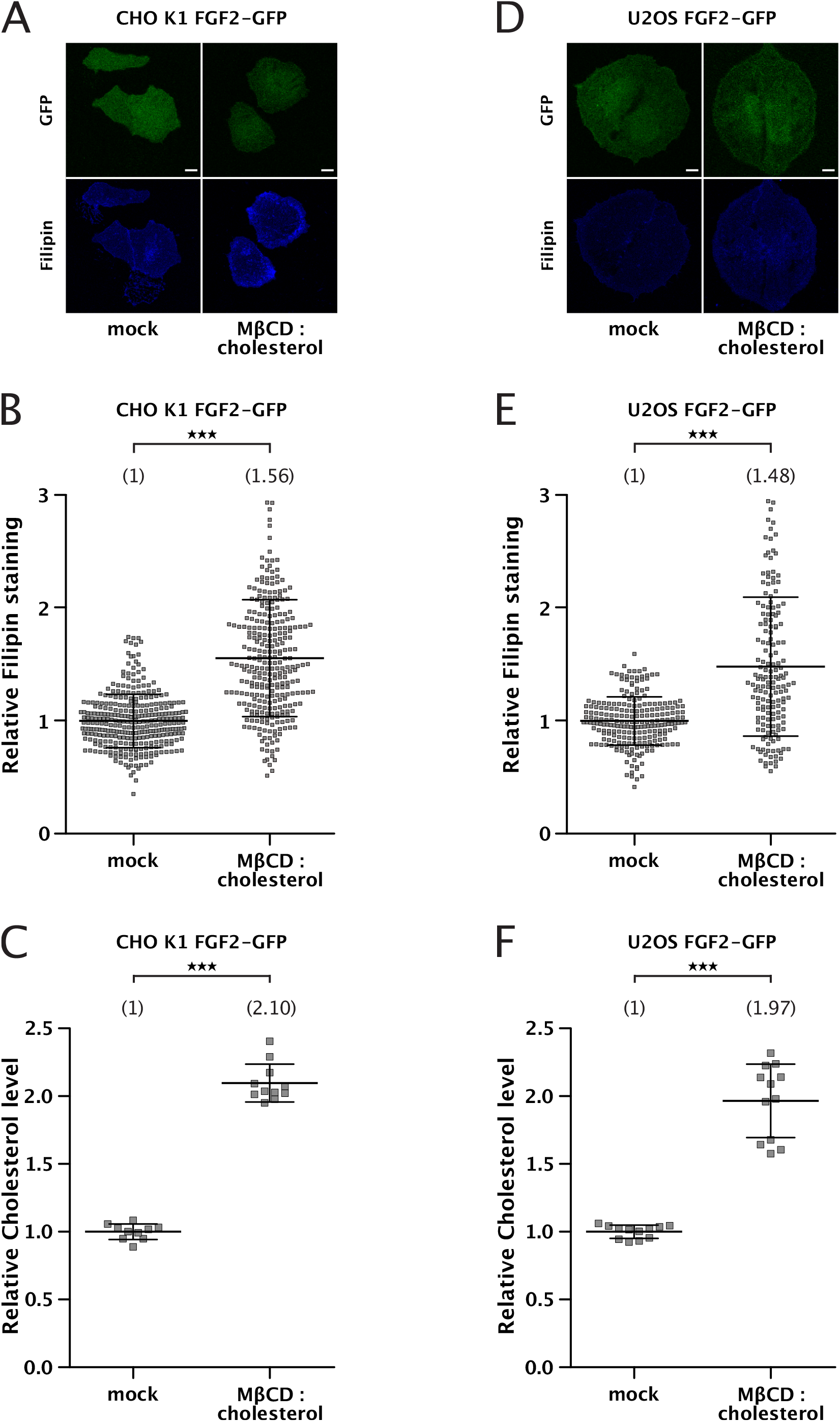
Quantification of cellular cholesterol levels using filipin imaging and mass spectrometry. Stable CHO K1 (**A**, **B**, **C**) and U2OS (**D, E, F**) cell lines were treated with Cholesterol:Methyl-β-Cyclodextrin (1:10 molar ratio) complexes for 1 hour in culture conditions. Cells were stained with filipin to visualize cholesterol using confocal microscopy (**A**, **B**, **D**, **E**). For each condition, both GFP and filipin fluorescence is shown (**A**, **D**; Scale bar = 6 μm). Confocal images were analyzed using ImageJ with the quantification of cholesterol levels for CHO K1 and U2OS for all conditions shown in panel **B** and **F**, respectively. The mean intensity values of the filipin signal detected per cell for each condition are shown in brackets, with the mock condition set to 1. Data are shown as mean ± SD (n = 4). The statistical analysis was based on a one-way ANOVA test (***p ≤ 0.001). In panel **C** and **F**, quantification of cholesterol levels using mass spectrometry is shown for CHO K1 and U2OS, respectively. Ratios of cholesterol to PC were determined for all different conditions indicated with the mock condition set to 1 for each cell line. Data are shown as mean ± SD (n = 4). The statistical analysis was based on a one-way ANOVA test (***p ≤ 0.001).

